# Molecular characterization of beast’s maguey (*Agave antillarum* Descourt), in the Dominican Republic

**DOI:** 10.1101/344259

**Authors:** José R. Núñez, Ineko Hoda

## Abstract

**Background:** This research project was proposed to determine the genetic variability (if any) of beast maguey (*Agave antillanarum* Decourt) populations, by means of molecular markers of the AFLPs (Amplified Fragment Length Polymorphism) type. For the determination of genetic variability, samples were collected from wild populations of maguey growing in some regions of the Dominican Republic, mainly the north, northwest, south and southwest regions, using statistical techniques already established for these cases. These samples were processed and analyzed in the plant molecular biology laboratory of the Centtro de Biotecnología Vegetal (CEBIVE) of Instituto de Innovación en Biotecnología e Industria (IIBI) in Santo Domingo, Dominican Republic.

**Results:** As for the genetic diversity is concerned, it was found that several of the populations studied differ from each other without taking into account the geographic distance where they grow. We concluded that there is a possibility that more than one species of *Agave antillarun* could be found in the wild in the Dominican Republic.

## 1. Introduction

### 1.1 Agave antillarum

*Agave antillarum* belongs to the Agavaceae family which are endemic in the Americas and the Caribbean being its greatest diversity in Mexico, this family includes 300 species and several genres, among which stands out the Agave. Species of Agave in general contain fibers that some are of great commercial value [1]. Native species have not been exploited yet for this purpose, but with the aim of curing diseases in man and animals by their great medicinal properties Use the “Insert Citation” button to add citations to this document.

The Agavaceae are particularly abundant in serofilos thickets, Chaparral, seasonally dry forests and deciduous tropical forests, there are species in any biome [2].

Although there are no reports of the genetic characteristics of the species of Agave in the Dominican Republic and their molecular characterization, the reports on these issues are abundant in the scientific literature for many species of this genus that grow in several areas of the world [3,4,5,6,7,8,9,10, 11, 12, 13].

The Agavaceae family are divided into the subfamilies Yuccoideae and Agavoideae, this last with approximately 190 species in about 8 genera. The genus Agave can be separated into two different subgenera, *Littaea* with racemose or spiked inflorescences and inflorescences and *Agave* with inflorescences in panicles branches or in umbels. The latter is distributed in the tropical and sub-tropical areas of the world and represents a large group of succulent plants with about 130 described species [1]. The genus *Agave* L (*Agavaceae*) L has its center of origin and diversification in Mexico, both in agriculture and in the biological sense, where spread throughout Mesoamerica and the Caribbean [8].

The beast maguey is an endemic species that grows in different regions of the country, especially in those places where there is a shortage of rainfall and low fertility soils [14]. Since Indians inhabited the island of Hispaniola, mixed-use have been given to the different species of Maguey, attributing them medicinal properties, insecticidal and as food [7]. Since this is a species that is threatened by indiscriminate collection and overgrazing by goats, it is necessary that this genetic resource be characterized molecularly, be conserved.

### 1.2 Genetic variability

Knowledge of the genetic diversity within and among plant populations, as well as the type of variation that leads to the formation of new species, is an essential part of the studies on evolution [15]. Many of the Agave species are of vegetative reproduction via rhizomes that grow from the base of the plant or by bulbils that originate in the floral peduncle. The plants formed by any of the two forms should be identical clones of the mother [16], however, other species of this genus produce viable seeds. The genetic diversity of the species of *Agave* of the Dominican Republic is unknown but the fact that many species are cross-pollinated, suggests that there may be some genetic variability in populations of *A*. *antillarum* from our country [8].

### 1.3 Medicinal properties of the *Agave*

Many species of Agave are of great economic importance such as henequen (*Agave fourcroydes* Lem.) and sisal (*Agave sisalana Perr*.) from which hard and strong fibers are extracted for making ropes, bags and rugs. Other species are used for the manufacture of alcoholic beverages such as tequila (*Agave tequilana* Weber) and bacanora (*Agave angustifolia* Haw.). some, like *Agave victoria-reginae* are prized as ornamentals especially for dry landscape. In addition, many of these plants are important raw materials for the industry sources such as steroids for pharmaceutical products, syrup of high content of fructose and inulin for the production of pre-biotics [17].

Inulin is a generic term for a polydisperse chain of fructose units who have degrees of polymerization, which vary from 2-60, usually with an average of ~ 12. The fructans are oligo - or polysaccharides comprising at least two monomers of adjacent fructose. They have value in health and occur in nature in a polydisperse way. The degree of polymerization has to do with the functional behavior of fructans and determines their end use. Molecules with a degree of polymerization of 2-7 are known as oligofructose, while larger molecules are known as inulin [2].

The genus Agave that grows in the Dominican Republic has not been studied in detail to determine the chemical components with pharmacological and food properties and its content of these chemicals is not known today.

*Agave* spp. are widely used in all rural areas of the Dominican Republic. They have been used in the treatment against cancer of bile bladder, ulcers, tumors, inflammation, high blood pressure, uterine fibroids, as well as an anti-tuberculosis agent. The water from the roots is used as a laxative and leaves crushed with salt and rum is applied to ulcers of domestic animals, hence their vernacular name (maguey de bestia). The inner parts of the leaves are applied over the belly to induce evacuation of the bladder and bowels [14].

## 2. Materials and methods

### 2.1. Sampling

The samples for this study were collected in regions of the country where this species is found in the wild. These regions are as follows: North (La Vega, Santiago and Puerto Plata); Northwest (Santiago Rodriguez, Valverde, Monte Cristi and Dajabón); Sur (San Cristóbal, Peravia, San José de Ocoa, Azua and San Juan de la Maguana); and Southwest (Bahoruco, Barahona, Pedernales, independencia). A number of samples were taken of each region first constructing a sampling frame for the distribution of the populations in the country making voyages of exploration to identify the places where there are significant concentrations of beast’s maguey. The number of samples to take was according to the existing population. This sample consisted of young plants. Sampling sites were of approximately 500 m^2^ with a separation between site of at least 250 m. 10 plants were collected by sampling site according to the methodology described by [15]. The sampling was carried out from a meaningful summary of the total universe of stocks for each zone in particular, depending on the number of plants existing in that area. It was collected material only from the above-mentioned sources. The collected material was properly identified and transported to the CEBIVE where prepared material for in-vitro planting and sowing in the field for later use.

### 2.2. Extraction, purification and quantification of DNA

The following general protocol published by Nickren [18], which was modified and adapted for Agave in our lab, was used for extraction and purification of DNA from the beast’s maguey leaves.

Plant Genomic DNA Extraction using CTAB: Large-Scale

Adapted from Daniel L. Nickren: Molecular Methods in Plant Biology, Fourth Edition, 2006. Department of Plant Biology Southern Illinois University

Before you begin, turn the centrifuge for pre - cool camera. The rotor is normally saved pre-cooling. It is assumed that all glass and plastic utensils are clean and sterile (autoclaved). You will need the following:

##### Equipment

- Centrifuge (with rotor)
- Polypropylene centrifuge tubes (30 and 50 ml capacity)
- Cheese cloth (cut into squares of 6 “x 6”)
- Funnels
- Heater with beaker and water
- Thermometer
- Liquid nitrogen
- Water bath to 37 - 45 ° C (JULABO USA, Inc., Allentown, PA)

##### Reagents

- Buffer CTAB 2X: 100 mM Tris-HCl, 1.4 M NaCl, 30 mM EDTA, 2% (w/v) CTAB, (0.1% N-Lauroylsarcosine and 1% PVP). The buffer is usually at pH 8.0 without adjusting.
- Protease K (Sigma Type H, at 0.13 units/mg, P-4755). You can save this enzyme in aliquots of 50 units/ml in the refrigerator.
- Dithiotreitol (DTT) 0.5 M. Store in aliquots of 0.5 ml - add to the buffer just before use.
- Buffer TE 1X
- **Ammonium acetate, 4.0 M.**
- **Organics:** chloroform: isoamyl alcohol (24:1), equilibrated phenol with Tris, cold isopropanol, cold 100% ethanol and cold 70% ethanol.
- **2-**Mercaptoethanol.

The best extraction of DNA is obtained from young plant material. Try the extraction as soon as possible after collection, if it cannot be chilled. For this procedure you will need about 3 g of plant material.

##### Protocol

1. Het a water bath in a heater just below boiling (95 ° C). Add 25 ml of CTAB buffer and 25 µl of 2-Mercaptoethanol to the tubes.
2. Place 2 to 3 g of plant material in the mortar, cover it with liquid nitrogen and grind it to a fine powder.
3. Add 25 ml of warm CTAB buffer and continue grinding for one minute more. Using a funnel with cheesecloth, squeeze the extract into a polypropylene centrifuge tube.
4. Add 50 units of protease K and 0.5 ml of DTT, mix, cover and incubate sample in a water bath at 37 - 45 ° C for an hour with occasional agitation.
5. Add 2/3 volume of chloroform: isoamyl alcohol (24:1). CAP the tube and shake gently for 2 to 3 minutes, intermittently releasing pressure.
6. Balance tubes (one with the other or with a blank one) using a plate scale. All the subsequent stages of centrifugation are assumed that the tubes are balanced. Centrifuge at 8,500 rpm for 15 minutes.
7. Remove the upper (aqueous) phase with a wide-tipped pipette) and place it in a clean polypropylene tube. **Be careful not to include anything of chloroform or interface.**
8. Add 2/3 volume of cold isopropanol and place it in a refrigerator to - 21 ° C for at least one hour (overnight is OK).
9. Centrifuge for 20 minutes at 10,000 rpm.
10. Carefully remove the supernatant, leaving the precipitate in the bottom of the tube. Invert the tube over a tissues paper towel to drain excess of isopropanol. If the precipitate starts to slip down, place tube upside up. Air dry the precipitate until it is slightly moist (at least one hour).
11. Add 3.0 ml of TE and 2.0 ml of 4.0 M NH4OAc. Using a 1.0 ml pipette tip to which the final part has been cut, suspend the precipitate in the buffer. It is advisable not to pipette very strong to not break the DNA.

***Next step phenol/chloroform is optional, but helps eliminate any protein that remain, only continue with precipitation of ethanol in step 13.***

12. Add 3.0 ml of phenol (saturated with Tris, pH 8.0 - bottom phase! And 3.0 ml of chloroform. Shake for one minute and centrifuge at 8,500 rpm for 15 minutes. Remove the upper aqueous phase and place it in a centrifuge tube. Re-extract with an equal volume of chloroform. Centrifuge at 8,500 rpm for 10 minutes. Remove the aqueous phase to a new centrifuge tube.
13. Add 2.0 volume of 100% ethanol. Store at - 20 ° C for at least 1 hour.
14. Centrifuge at 10,000 rpm for 20 minutes. Remove the ethanol, rinse briefly with 70% ethanol and dry the precipitate DNA investing on a washcloth tissues. Allow the DNA to dry completely at room temperature (overnight).
15. Re-suspend the DNA in TE, one ml if the precipitate is large or 0.5 ml if it is small. Use a cut pipette tip to place the DNA in a 1.5 ml tube of 1.5 ml labeled.
16. If you want to, RNA can be eliminated from the sample of DNA by digestion with RNase. You can use a combination of two enzymes, RNase A (at 10-50 µg/ml of sample) and RNase T1 (at 50 units/ml of sample). Incubate the samples for 30 minutes at 37 ° C. Do a phenol/chloroform extraction and then a chloroform extraction as indicated in step 12 above. Precipitate DNA with ethanol as indicated in steps 13-15.

### 2.3. Quantification of DNA

DNA quantification was performed using a spectrophotometer SHIMADZU BioSpec-nano brand, Shimadzu Corporation, Kyoto, Japan. The apparatus was set to a pathlength of 0.2 mm for dsDNA to determine from 50 to 3,700 ng/µL.

## 3. **Results and Discussion**

### 3.1. AFLP Determination

AFLP^TM^ (amplified fragment length polymorphism) technique is used to display hundreds of amplified DNA restriction fragments simultaneously. The AFLP banding patterns, or fingerprints, can be used for many purposes, such as monitoring of the identity of an individual or the degree of similarity between isolated individuals. Polymorphisms in the patterns of bands map specific loci, allowing genotyped individuals or differentiated based on the basis of the alleles carrying (AFLP^®^ Plant Mapping Protocol, Applied Biosystems^®^, 2010).

AFLP technology combines the power of Restriction Fragment Length Polymorphism (RFLP) with the flexibility of the technology based on PCR linking sequences recognized by the primers (adapters) to DNA restriction. (For more details on) this topic please consult the document cited above in: http://www3.appliedbiosystems.com/cms/groups/mcb_support/documents/generaldocuments/cms_040959.pdf

For the determination of the AFLPs, the kit named “AFLP Ligation and Preselective Amplification Module for Regular Plant Genome (500-6000 Mb)” from Applied Biosystems^®^ was used (module of ligation and Preselectiva amplification of AFLP for plants genome [Regular] 500-6000 mb]) P/N: 402004. This kit contains EcoRI adapter, adapter MseI, the pre-selective amplification primers, preselective amplification, main mix (Amplification Core Mix) and control DNA. The enzymes (EcoRI, MesI, and T4 DNA ligase) were obtained from New England BioLabs^®^ Inc. 240 County Road Ipswich, MA 01938-2723. Reagents for the genetic analyzer, (Hi-Di Formamide, Pop7 Polymer, GeneScan 500 Buffer) were acquired from Life Technology^®^, Carlsbad, CA, United States of America, subsidiary company of Applied Biosystems^®^).

Five hundred (500) ng of genomic DNA were digested with the restriction enzymes EcoRI/MseI. Restriction. amplification of the digested fragments was carried out using a system of fluorescent dyes supplied by Applied Biosystems^®^) in the kit called “AFLP^TM^ Plant Mapping”. This system is ready to be used with the genetic analyzer ABI PRISM^®^ 3130xl. This software has the ability to identify common polymorphic peaks among a large number of samples. The AFLP analyses were made following instructions provided by the suppliers of the analysis kit. Linkage and AFLP Amplification modules were purchased directly from the manufacturer through its supplier in Puerto Rico (BioAnalytical Instruments PO BOX 270021, San Juan, Puerto Rico 00927). Three different fluorescent dyes were used in the analysis of AFLP to determine if there was a genetic diversity in the populations of beast’s maguey in the Dominican Republic. These dyes were FAM (5’ 6-FAM (Fluorescein)) and JOE (6-JOE, SE (6-Carboxy-4’,5’-Dicloro-2’,7’-Dimetoxyfluorescein, Succinimidil Ester).

The analysis of the results of the running in the automatic genetic analyzer were made using the software “GeneMarker^®^” from SOFTGENETICS^®^, State College PA 16803 USA. The dendrograms and statistics were generated using the free access programs

PowerMarker v3.0 (http://www.powermarker.net),

TreeView (http://taxonomy.zoology.gla.ac.uk/rod/treeview.html),

TreeGraph 2 v2.4 (http://treegraph.bioinfweb.info/)

and FigTree (http://tree.bio.ed.ac.uk/software/figtree/).

### 3.2. Determination of AFLP using the fluorescent dye FAM

This dye (blue) was used applying the combination of primers EcoRI-ACA-MseI-CTG. The combination Also tested the EcoRI-ACA-MseI-CAG combination with good results, but these were less consistent than the first combination. EcoRI-ACA-MseI-CAG was also tested with good results but these were less consistent than the first combination

The correlation coefficient for the different populations in this type of analysis is shown in Figure 3.2.4. Blue colored cells show population differences were found with the analysis of conglomerate of GeneMarker^®^ program mentioned above. This correlation analysis shows difference between the populations of Bahoruco and Azua. Azua and Puerto Plata; Azua and San Cristobal; Independencia and Bahoruco; Independencia and Puerto Plata. You can notice that the genetic difference is quite independent of geographical distance of the populations although the geographic distance affects enough genetic difference as it is the case of San Jose de Ocoa-Bahoruco ; San Jose de Ocoa - Puerto Plata; Mao-Azua; Barahona, Puerto Plata; La Vega (Constance) - Puerto Plata; Azua-Santiago Rodríguez and other populations that show differences as the geographical distance is lengthened.

**Figure 3.2.1.**
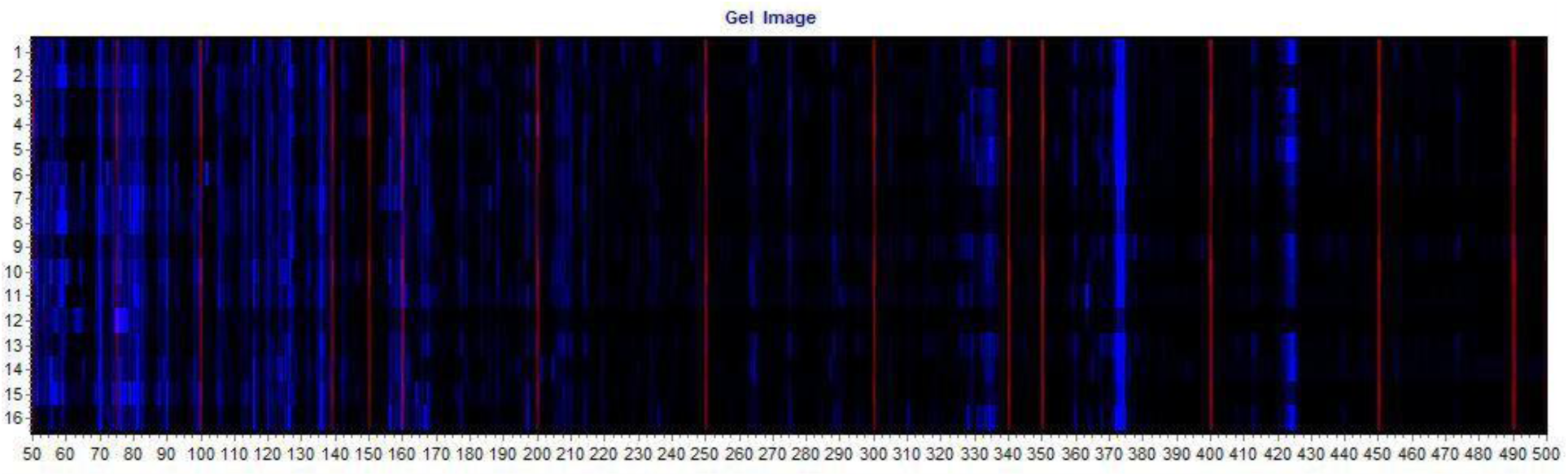
Representation of the gel obtained for the run with the combination of EcoRI-ACA-MseI-CTG primers. The Red strips represent the coloring standard (GeneScan-500).

**Figure 3.2.2.**
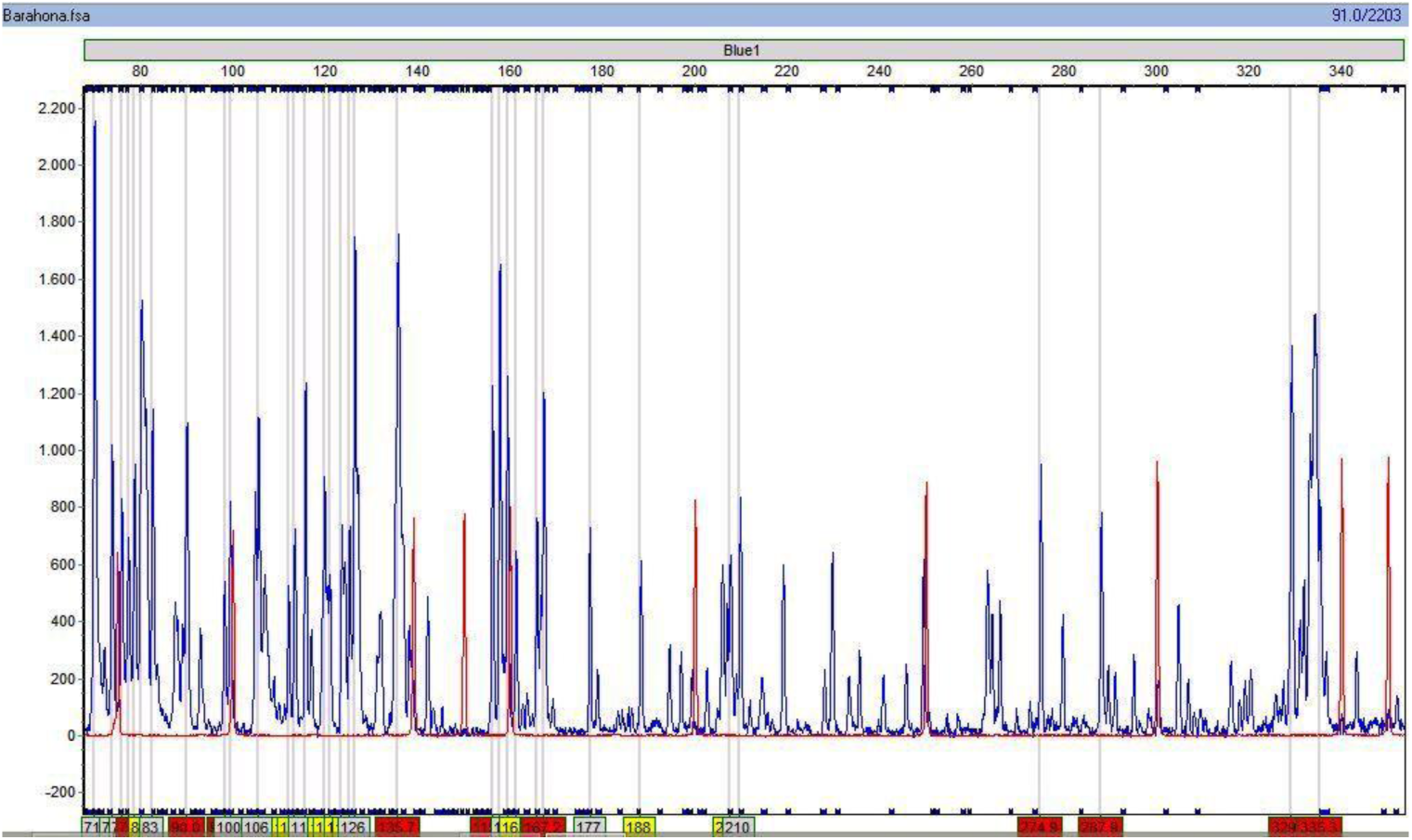
Representation of the peaks obtained from one of the samples analyzed in the automatic sequencer ABI PRISM^®^ 3130xl. The Red peaks are coloring standard GeneScan-500. The grey bands are the tlocations (bin) established for the peaks.

**Figure 3.2.3.**
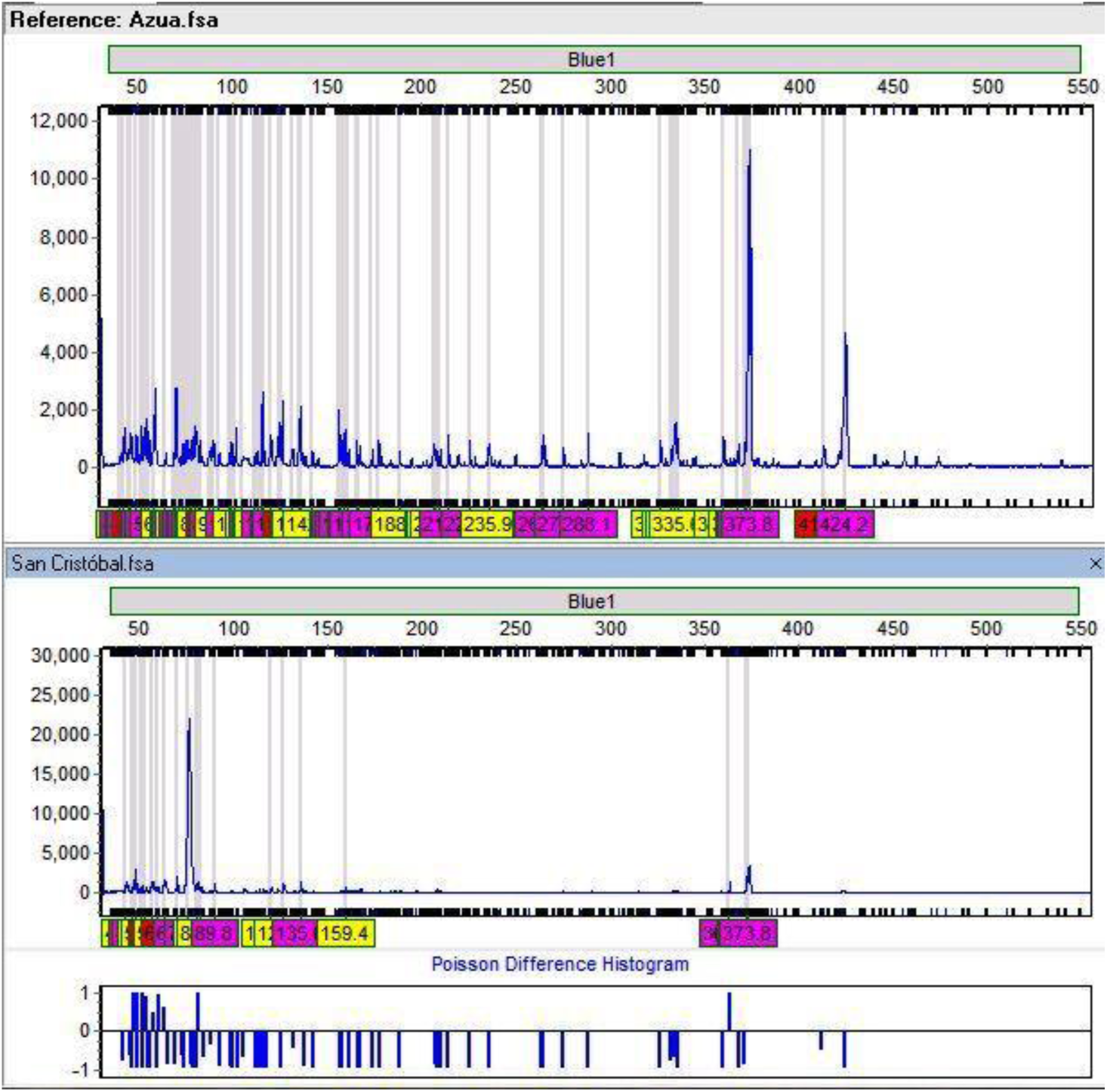
Poisson difference between beast maguey populations that grow wild in Azua and San Cristobal. Note the absence of peaks in the population of San Cristobal that are present in the population of Azua.

**Figure 3.2.4.**
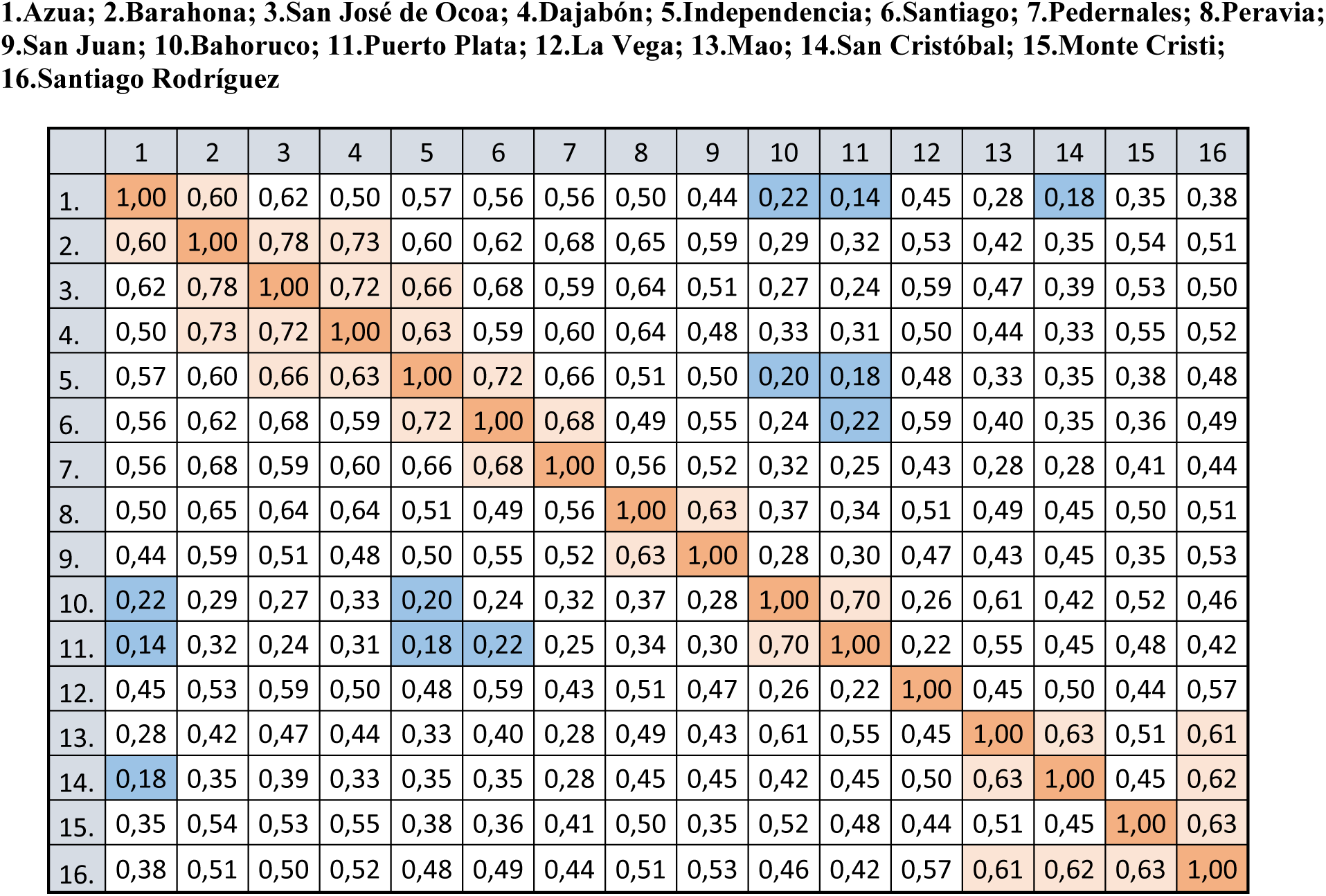
Correlation coefficient for ACA-CTG (FAM)

The dendrogram shown in Figure 3.2.5 derives from this correlation and the same three groups or clusters can be distinguished. Group I consist of the populations of Azua, Barahona, San José de Ocoa, Dajabón, independencia, Santiago, Pedernales, Peravia and San Juan; the components of Group II are Bahoruco and Puerto Plata, while group III are Mao, San Cristóbal, Monte Cristi and Santiago Rodríguez. La Vega is isolated as a lone population though quite related with the other populations with the exception of Bahoruco and Puerto Plata.

**Figure 3.2.5.**
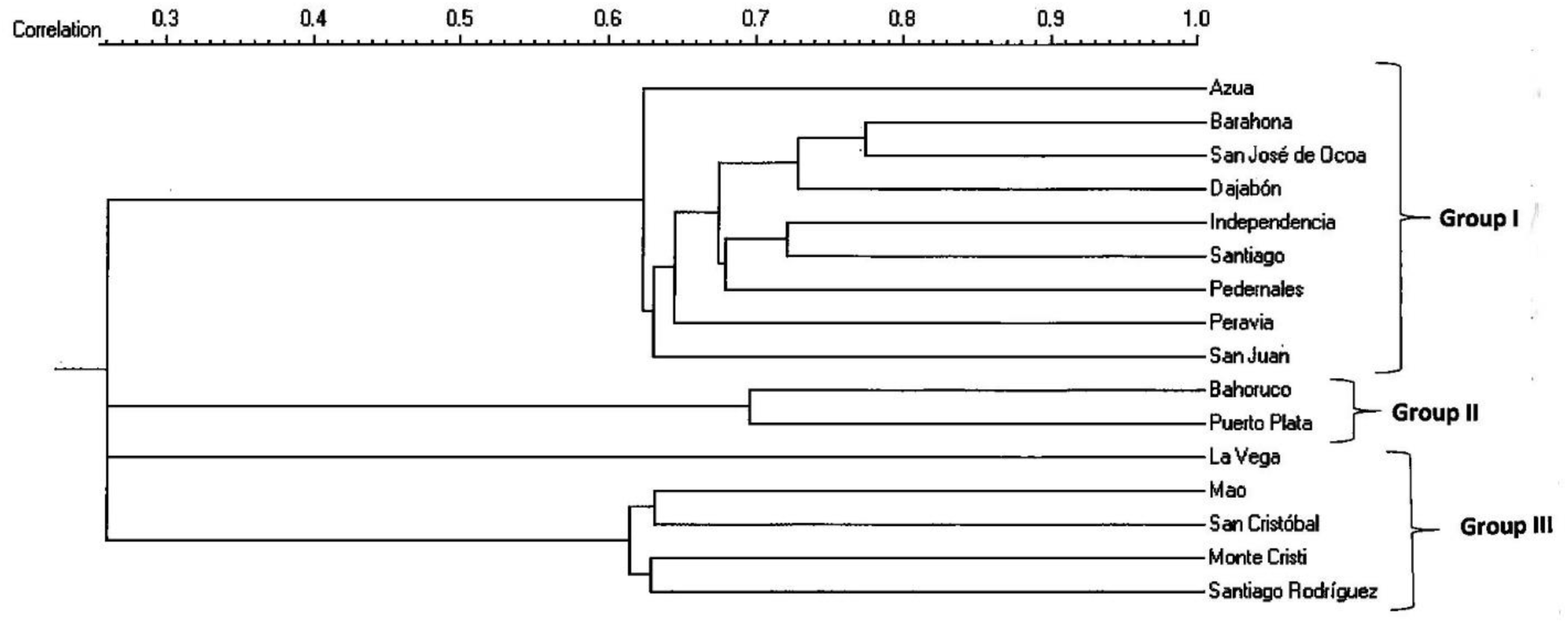
Dendrogram generated with data from the coefficient of correlation of different populations of wild beast’s maguey of the Dominican Republic using dye FAM and the combination of EcoRI-ACA-MseI-CTG primers.

**Figure 3.2.6.**
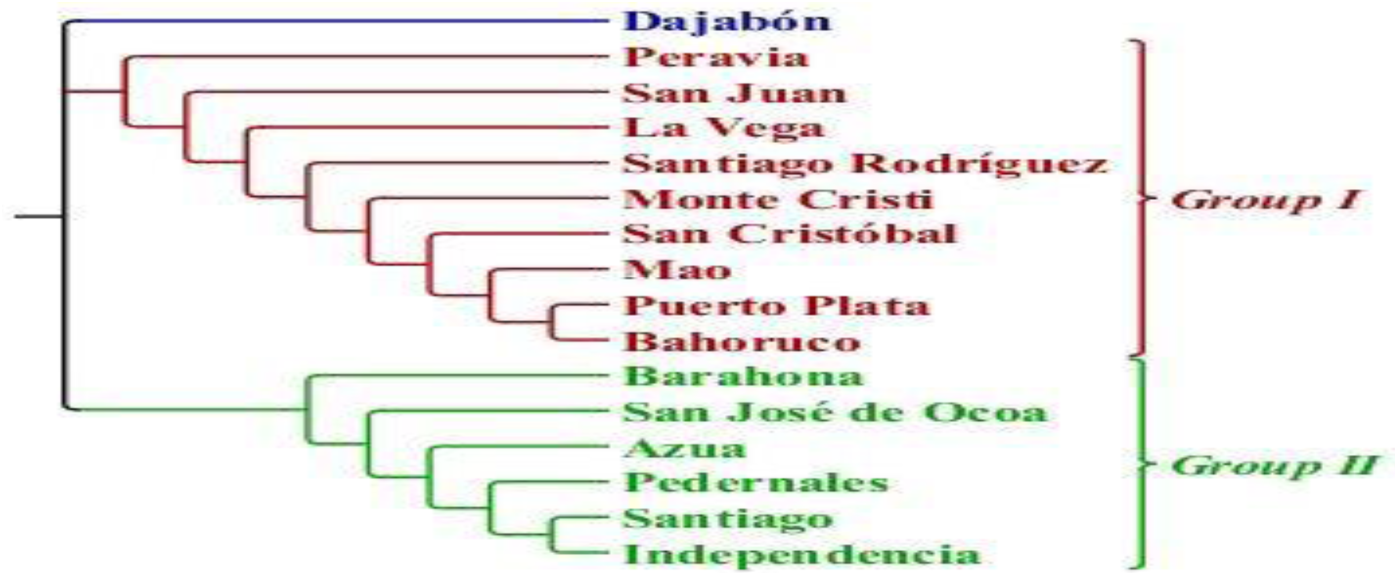
Dendrogram generated by “bootstrap Neighbor Joining” with the data obtained by the GeneMarker^®^ program. Data of the different populations of wild beast’s maguey from the Dominican Republic using the dye FAM and the combination of EcoRI-ACA-MseI-CTG primers.

With the data obtained with the GeneMarker^®^ program, a matrix was created to run the program PowerMarker v3.0 with which a “bootstrap” was applied to the data and thus a dendrogram was generated with Euclidean distance, which is shown in Figure 3.6.

Compared to the correlation dendrogram, this dendrogram shows only two groups and an isolated population (Dajabón). Group I consist of Peravia, San Juan, La Vega, Santiago Rodríguez, Monte Cristi, Puerto Plata, San Cristóbal, Mao and Bahoruco, while group II is formed by Barahona, San José de Ocoa, Azua, Pedernales, Santiago, and independencia.

The order of the groups is inverted in this dendrogram in comparison with the previous one. However, all components of Group II are included in Group I of the correlation coefficient dendrogram indicating the same relationship of the populations. Puerto Plata and Bahoruco, which form Group II of the former dendrogram and which are found in group I of the “bootstrap” dendrogram, show the same similarity in the analysis using both systems. The components of Group III of the correlation coefficient dendrogram (Mao, San Cristóbal, Monte Cristi and Santiago Rodriguez) are included in Group I of the “bootstrap” dendrogram. We can deduce, by analyzing this dendrogram, that the differences among populations analyzed with the system “bootstrap” are the same as those found with correlation analysis.

### 3.3 Determination of AFLP using the fluorescent dye JOE

This (green) dye was applied using the combination of EcoRI-AGG-MseI-CAA primers. The representation of the gel of the run with this dye and combination of primers is shown in Figure 3.3.1 below.

**Figure 3.3.1.**
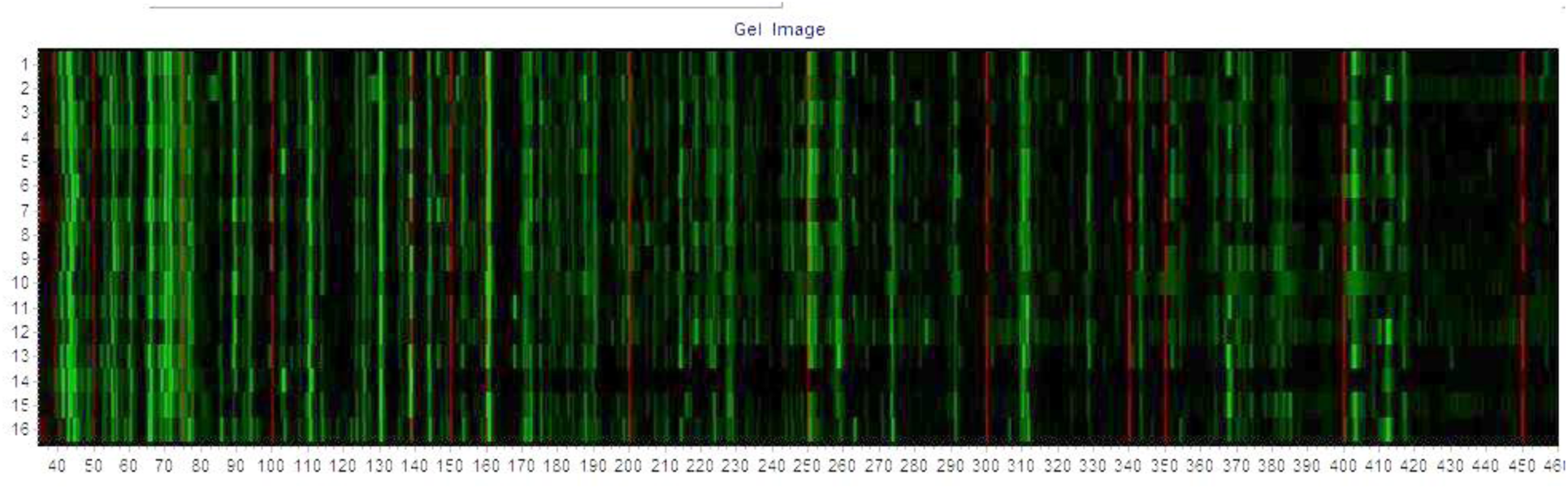
Representation of the gel obtained for the run with the combination of EcoRI-AGG-MseI-CAA primers. The Red strips represent the coloring standard (GeneScan-500).

A sample of the typical peaks of one of the runs of this system of dyes/primers are shown below in figure 3.3.2.

**Figure 3.3.2.**
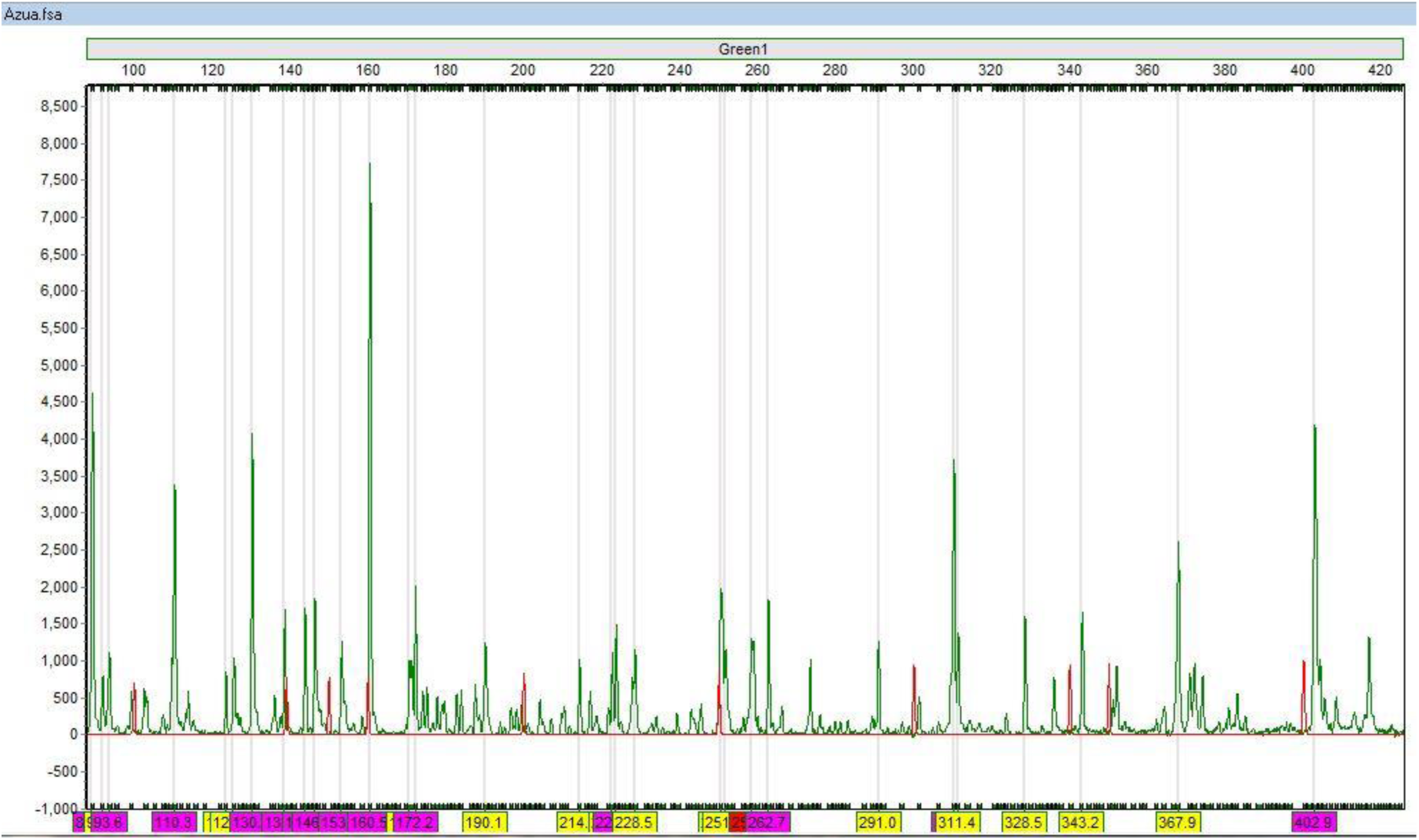
representation of the peaks obtained from one of the samples analyzed in the automatic analyzer ABI PRISM^®^ 3130xl with the combination of the dye JOE and EcoRI-AGG-MseI-CAA primers. The Red peaks are of the coloring standard GeneScan-500. The grey bands are the locations (bin) established for the peaks.

Using this combination of primers/dye (EcoRI-AGG-MseI-CAA (JOE) genetic difference between some of the populations of beast’s maguey was found as can be seen in the analysis of fragments comparison and the Poisson difference displayed in figure 3.3.3. In this figure you can see the genetic differences between the populations of Azua and San Cristobal.

**Figure 3.3.3.**
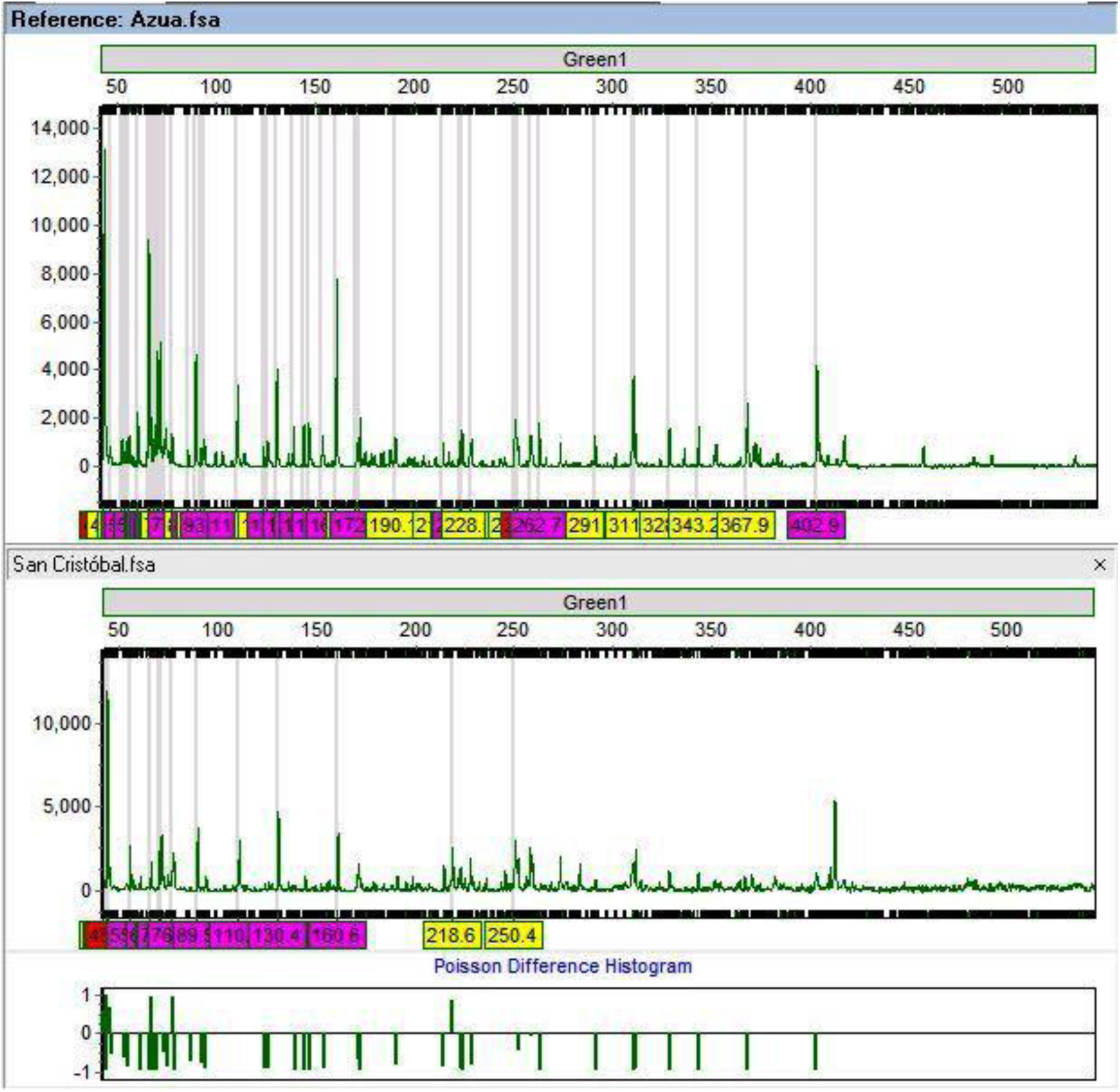
Poisson difference between beast’s maguey populations that grow wild in Azua and San Cristobal. Note the absence of peaks in the population of San Cristobal that are present in the population of Azua.

The correlation coefficient for the different populations in this type of analysis is shown in Figure 3.3.4. Blue colored cells show population differences found with the cluster analysis of the GeneMarker^®^ program mentioned above. This correlation analysis shows difference between the populations of Bahoruco and Azua. Azua and San Cristobal; Santiago and Bahoruco; Santiago and San Cristobal; San Cristóbal and Mao, among others. Here you may also notice that the genetic distance between populations is quite independent of geographical distance but the biggest genetic difference found in distant populations with the exception of Azua-San Christopher.

**Figure 3.3.4.**
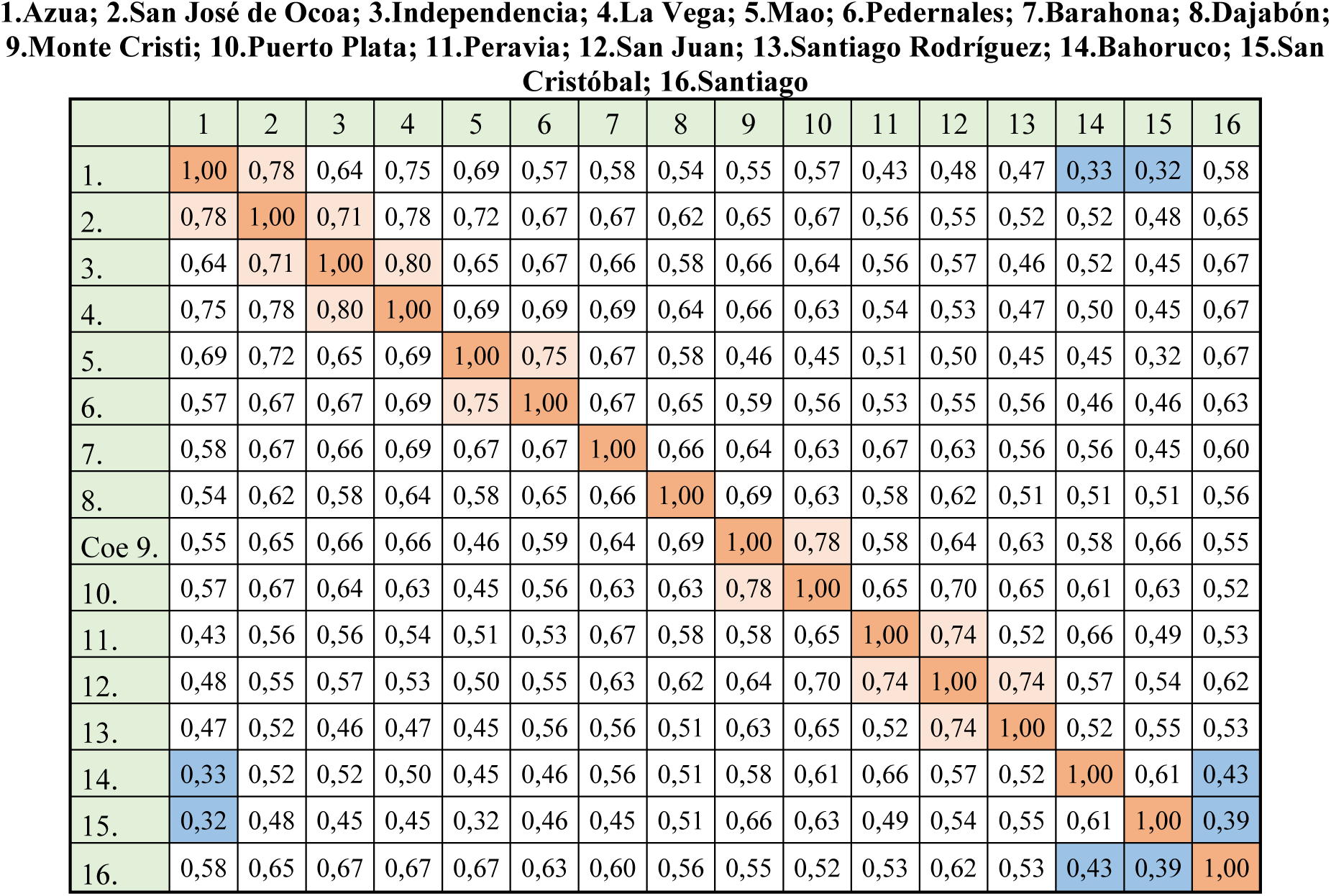
Correlation Coefficient for AGG-CAA (JOE)

The dendrogram shown in Figure 3.3.5 derives from this correlation and on it you can see a large group or cluster and three isolated populations (Bahoruco San Cristóbal and Santiago). Group I can be subdivided into two subgroups. The subgroup Ia consists of the populations of Azua, San José de Ocoa, independencia, La Vega, Mao, Pedernales and Barahona. The subgroup Ib is composed of the populations of Dajabón, Monte Cristi, Puerto Plata, Peravia, San Juan and Santiago Rodríguez.

**Figure 3.3.5.**
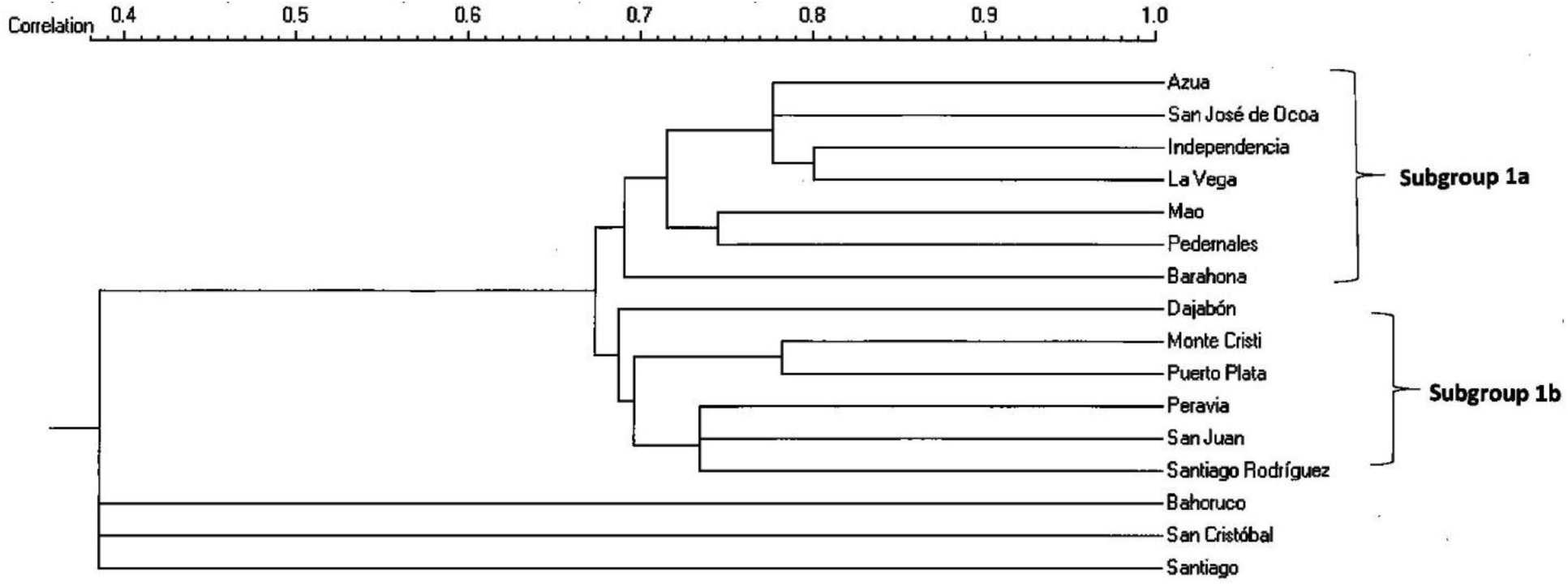
Dendrogram generated with data from the correlation coefficient of the different populations of wild beast’s maguey from the Dominican Republic using the dye JOE and the combination of EcoRI-AGG-MseI-CAA primers.

With the data generated with the GeneMarker^®^ program, a matrix was developed that was run in the PowerMarker program v3.0 with which a “bootstrap” was applied to the data to generate a dendrogram with Euclidean distance, which is shown in figure 3.3.6.

**Figure 3.3.6.**
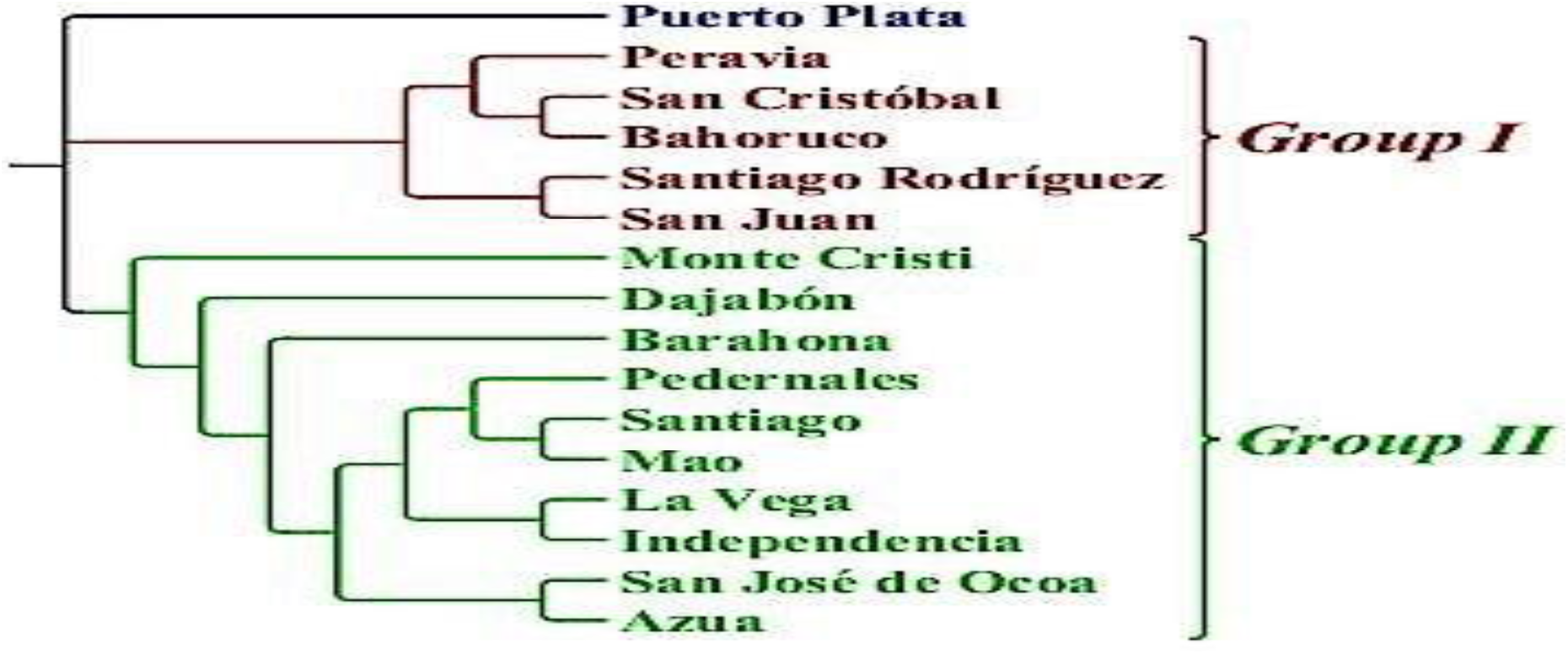
Dendrogram generated by “bootstrap Neighbor Joining” with the data obtained by the GeneMarker^®^ program. Data of different populations of wild beast’s maguey of the Dominican Republic using the coloring JOE and the combination of EcoRI-AGG-MseI-CAA primers.

Compared to the dendrogram of correlation, this dendrogram also presents two groups with an isolated population (Puerto Plata).

The majority (70%) of this dendrogram Group II populations are included in the subset I of the dendrogram of correlation, indicating good relationship of similarities. Only 60% of the populations of Group I are included in subgroup II of the correlation dendrogram, but it must be taken into consideration that these dendrograms were generated with two different algorithms (Euclidean distance and correlation coefficient) for which the differences in similarities tend to be marked.

However, the greatest differences among populations remain the same (Azua-Bahoruco; Azua-San Cristobal; Santiago-Bahoruco; (and Santiago-San Cristobal). It can be inferred, analyzing this dendrogram, that the differences among populations analyzed with the “bootstrap” system are the same as those found with the analysis of correlation coefficient.

## 4. Conclusions

The genetic variability of the beast’s maguey (*Agave antillarum*) can be seen by the phenotype in the initial growth phase of the plants from in vitro culture of some populations compared with others. In some populations (Santiago, for example) initial growth was always displayed with a purple/brownish coloration and more scattered thorns, while other populations (San José de Ocoa, for example, the initial growth always showed with a natural green color and with more numerous thorns, as shown in Figures 4.1a and 4.1b. This difference was less pronounced as the plants were more adult (see Figure 4.2 below)

**Fugure 4.1a.**
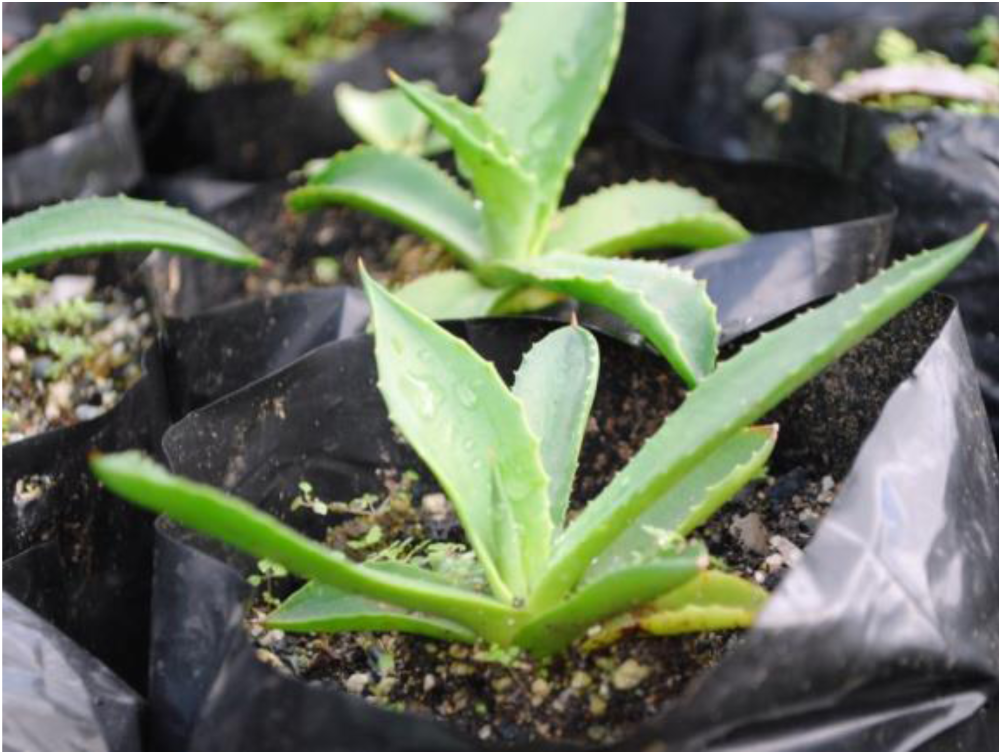
Population of San José de Ocoa.

**Figure 4.1b.**
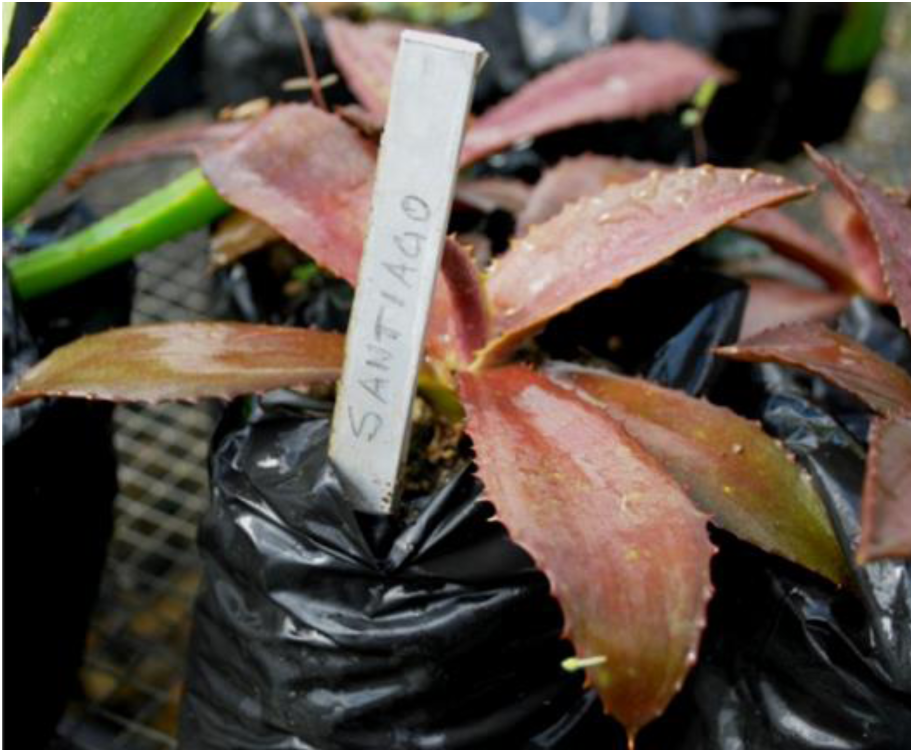
Population of Santiago.

**Figura 4.2.**
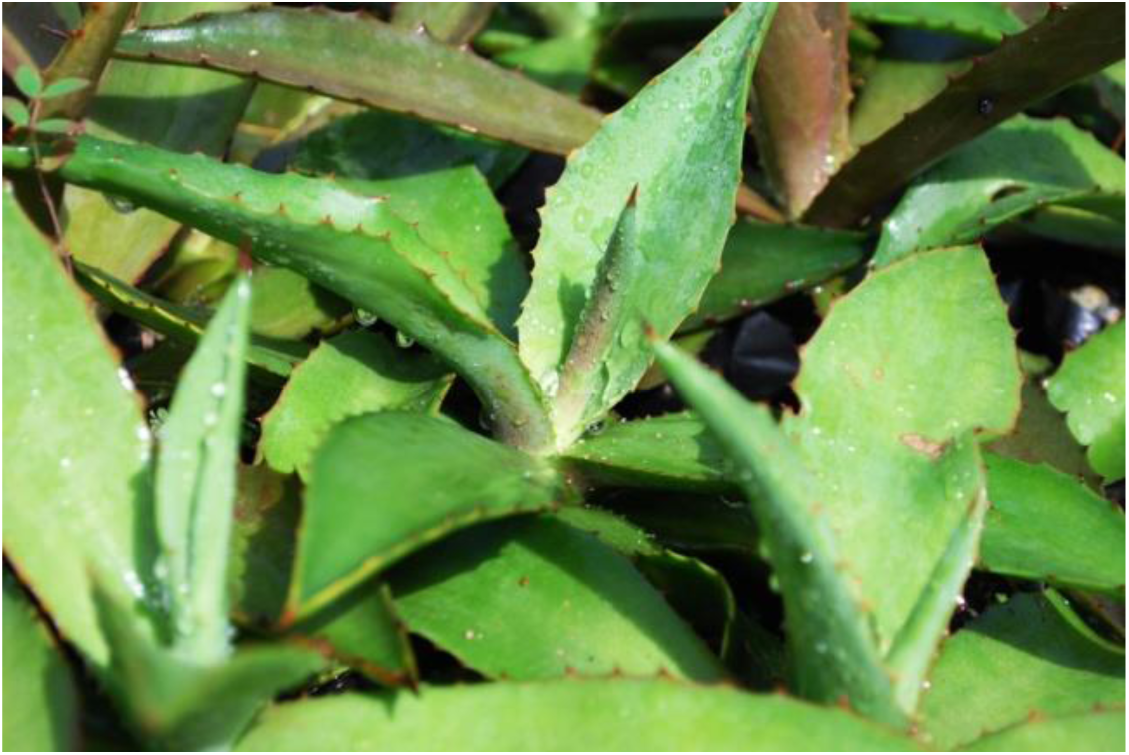
Planta de maguey de bestia de la población de Santiago más adulta. Nótese la coloración más pálida que la planta más joven de la Figura 5.1b.

The genetic variability was noted more markedly using AFLP technique to examine the population differences. Using the fluorescent dye FAM combined with EcoRI-ACA-MseI-CTG primers we found accentuated genetic difference between the populations of Azua and Puerto Plata; Azua and San Cristobal; Azua and Bahoruco; Puerto Plata and independencia; La Vega and Bahoruco; and La Vega and Puerto Plata.

Other populations showed less accentuated differences as they were cases of Ocoa-Bahoruco; Ocoa - Puerto Plata; Mao-Azua; Barahona, Puerto Plata; Puerto Plata-Pedernales; and Santiago Rodriguez-Azua.

Analyzing Figure 4.3 we can see that the geographical distance is not a main factor to make the genetic difference between populations, although it shows a tendency to the variability with increasing distance and geographic barriers that exist between populations (eg. Puerto Plata-Pedernales, 307 km; Puerto Plata-independence, 259 km).

**Figure 4.3.**
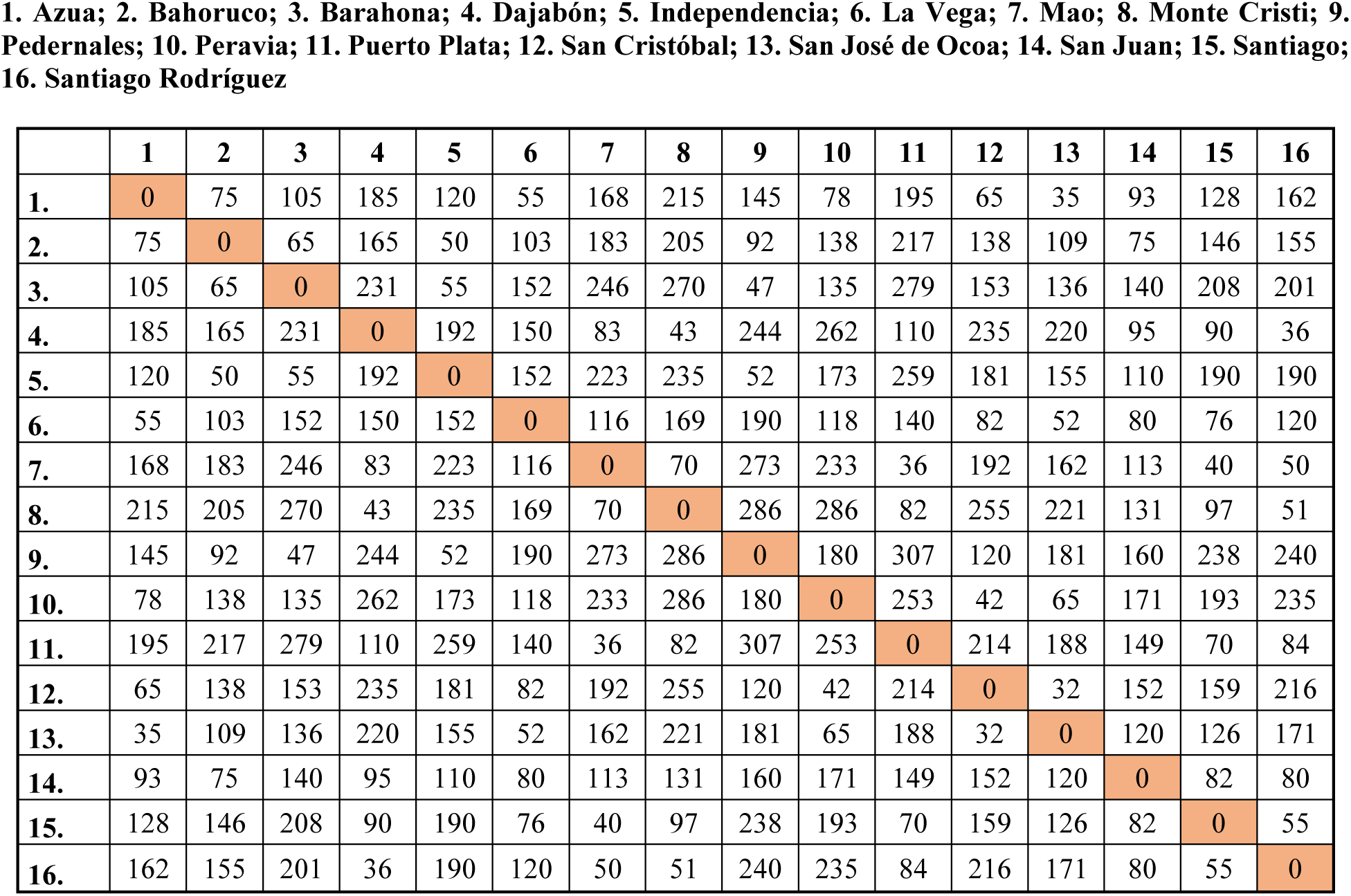
Geographical distance (Km) between wild populations of beast’s maguey from the Dominican Republic.

With the use of the fluorescent dye JOE in combination with EcoRI-AGG-MesI-CAA primers we also could find difference between populations, although not as clear as with the use of the fluorescent dye FAM. The correlation coefficient obtained with this dye (Figure 3.3.4) showed higher values than those obtained with the FAM dye (Figure 3.2.4). Four populations could be determined with variations between them. These were Azua-Bahoruco; Azua-San Cristobal; Santiago-Bahoruco; and Santiago-San-Cristobal. The populations Azua-Bahoruco and Azua-San Cristobal which markedly differ according to the data obtained with this dye/primer system also differ markedly with the data obtained with the FAM system indicating that there are real genetic differences between these populations.

It would be advisable a more detailed study of this species, both genetic and botanical, to determine the degree of variability that exists in between the different populations of beast’s maguey because as you could see, phenotypic variations can be noticed between the diverse populations in particular the population of Santiago, which showed a very peculiar feature in its early growth stage. The differences are so pronounced between this population and other populations that maybe we have a different subspecies of beast’s maguey in the Dominican Republic.

## Acknowledgements

Thanks to the area technicians of the Ministry of Agriculture who so kindly lent us their help on the journeys made across the country during sample collection. Thanks to technicians from the CEBIVE who helped us in collecting and processing samples. The Ministry of higher education, science and technology by the economic financing of this project and by the kindness of the extension of the deadline for the completion of this project. Any errors found in this report is the responsibility of the main author.

